# Validating the automation of different measures of high temperature tolerance of small terrestrial insects

**DOI:** 10.1101/2021.03.19.436121

**Authors:** Heidi J. MacLean, Jonas Hjort Hansen, Jesper G. Sørensen

## Abstract

Accurately phenotyping numerous test subjects is essential for most experimental research. Collecting such data can be tedious or time-consuming, and can be biased or limited by manual observations. The thermal tolerance of small ectotherms is a good example of this type of phenotypic data, and it is widely used to investigate thermal adaptation, acclimation capacity and climate change resilience of small ectotherms. Here, we present the results of automatically generated thermal tolerance data using motion tracking on video recordings using two *Drosophila* species and temperature acclimation to create variation in thermal tolerances and two different heat tolerance assays. We find similar effect sizes of acclimation and hardening responses between manual and automated approaches, but different absolute tolerance estimates. This discrepancy likely reflects both technical differences and the behavioral cessation of movement rather than physiological failure measured in other assays. We conclude that both methods generate biological meaningful results, which reflect different aspects of the thermal biology, find no evidence of inflated variance in the manually scored assays, but find that automation can increase throughput without compromising quality. Further we show that the method can be applied to a wide range of arthropod taxa. We suggest that our automated method is a useful example of through-put phenotyping, and suggest this approach might be applied to other tedious laboratory traits, such as desiccation or starvation tolerance, with similar benefits to through-put. However, the interpretation and potential comparison to results using different methodology rely on thorough validation of the assay and the involved biological mechanism.

## Introduction

Phenotypic data are used for everything from mapping out genotype-phenotype associations, to measuring selection responses, and evaluating local adaptation of natural populations. The generation of high-throughput molecular data (i.e. genome and transcriptome sequencing) is increasingly common but generating the corresponding phenotypic data is a much needed, but rate-limiting, step in many studies (see e.g. Mackay and Huang 2018). Furthermore, the scoring of phenotypes relies on individual observations, which risks measurement error (reduced reliability and repeatability), observation bias, and increased variance in such data sets (Lachin 2004, Hazell, Pedersen et al. 2008, Holman, Head et al. 2015). Thus, there is great potential for methods that can both increase the speed and number of samples processed and reduce the risk of biased, inaccurate or imprecise phenotyping.

Critical thermal limits (CTLs) of ectotherms are widely used physiological phenotypes in studies of physiology, macro-ecology, and evolutionary biology (Sørensen, Nielsen et al. 2005, Kellermann, Overgaard et al. 2012, Sunday, Bates et al. 2013). Methods used to assess thermal limits are diverse but generally apply static (time to knock-down, TKD) or dynamic (critical thermal maxima in ramping assays, CTmax) methods (reviews by Lutterschmidt and Hutchison 1997, Sinclair, Coello Alvarado et al. 2015). Exact methodology (e.g. knock-down temperature and ramping rate) affects the absolute estimates (see e.g. Jørgensen, Malte et al. 2019), yet these estimates often are treated as a “real value” that are comparable and can be applied directly in ecological/evolutionary models (Sunday, Bates et al. 2014). Traditionally, the endpoint recorded has been the onset of muscle spasm, the loss of righting response, the loss of movement (coordinated or complete), or death (reviewed by Lutterschmidt and Hutchison 1997) and all rely on an observer judgement in real time. For example, lower limits are typically measured by the loss of movement or the loss of righting response however, these measures might be very different from the lower lethal limit at which survival is compromised. As such, there are inherent conceptual and practical limits and biases in these data, including the number of individuals that can be scored (see e.g. Allen, Chown et al. 2016), inter- and intra-observer differences, block effects for different replicates and treatments (Castaneda, Calabria et al. 2012). Regardless of the exact method used, interpreting CTLs and the patterns thereof, clearly relies on an assumption that CTLs are measured in an accurate and precise manner and that they are relevant for the biological question (Santos, Castañeda et al. 2011, Sørensen, Kristensen et al. 2016, Kingsolver and Umbanhowar 2018, Jørgensen, Malte et al. 2019).

To date, advancing the quantity and quality of phenotypic scoring has been made using video recordings (Woods and Bonnecaze 2006, Hazell, Pedersen et al. 2008, MacLean, Higgins et al. 2016) and software that can track movement from such recording (Baatrup and Bayley 1993, Fisher, Rodriguez-Munoz et al. 2016, Thoen, Kloth et al. 2016, Soto-Padilla, Ruijsink et al.2018). Burton, Zeis et al. (2018) described a method for automated temperature tolerance of aquatic ectotherms, based on behavioral (movement) tracking of video recordings. Similarly, Awde, Fowler et al. (2020) described methods for scoring of cold knock-down (in a knock-down tube) and heat knock-down assays based on semi-automation. While all of the above these are examples of generating phenotypic data that rely less on real time observations, it is not clear how comparable these are to the traditional assays based on real time observations, i.e. if they lead to comparable estimates in relative or absolute terms. Moreover, it has not been established whether they provide more precise or accurate estimates of the endpoints as there have been no studies directly comparing to traditionally scored assays.

Here we evaluate a high-throughput automation, based on motion tracking of video recording, for the behavioral upper thermal limits in *Drosophila*. Specifically, we measure the behavioral CTmax (at different ramping rates and developmental acclimation temperatures), and behavioral TKD (at different temperatures and heat hardening treatments) in two species of *Drosophila* with known, markedly different, thermal tolerance and acclimation capacity (Sørensen, Giribets et al.2019) and compare to the most common method of measuring physiological thermal limits in *Drosophila* (Jørgensen, Malte et al. 2019). We expect that there will be some differences in the absolute values of the behavioral assay as the arena and thermal chambers are different between the two set-ups. Thus, successful automation as compared to manually scored methods were evaluated based on: 1) Species differences in thermal tolerance assays should be comparable, 2) Effect sizes of pre-treatments and treatment conditions should be comparable, 3) Automation should be high-throughput, 4) comparable absolute CTmax and TKD estimates from manual and automated assays, and 5) Variance attributable to observer imprecision should be reduced.

## Materials and methods

### Experimental animals

We used a mass bred populations of *Drosophila melanogaster* and *Drosophila subobscura* (established from 25 isofemale lines), collected from the Danish peninsula of Jutland in 2013 (Sørensen, Kristensen et al. 2015, Schou, Mouridsen et al. 2017). Flies were maintained under 12:12 h light/dark cycle at 19°C on a standard media (agar, oatmeal, yeast, sugar). Experimental animals developed under density-controlled conditions of 40 (± 5) eggs on fresh 7 mL food vials. Once emerged, flies were sexed and males were kept at a density of 30 individuals per 7 mL fresh food vial and allowed a minimum of 48 h to recover from anesthesia prior to experimentation.

### Experimental set-up

Manually scored assays for physiological thermal limit: a rack with randomly placed 5 mL screw-top glass vials each with a single male fly was placed in a water tank connected to a circulating programmable water bath (Julabo FP51-SL). A single observed scored all flies in a block, by observing flies until the defined endpoint, the cessation of all movement, was reached. Automated method for behavioral thermal limit: male flies were individually aspirated into experimental wells in 36-well plexiglass arenas (Rohde, Madsen et al. 2016) and placed on a lightbox in a temperature controlled cabinet (POL-EKO ILW 115 TOP+). Video recordings of the assays (filmed by a Samsung Tab A SM-T550 Tablet) were analyzed by Ethovision XT video tracking software (Noldus, Wageningen, The Netherlands) to generate activity over time. For both manual and automated assays the realized temperature in the waterbath/incubator and the experimental arena (either glass vial or plexiglass arena) were documented using dataloggers (see Results).

### Critical thermal maximum

CTmax was assessed using two ramping rates (targeted at 0.1 and 0.5 °C/min starting at 22.5°C) following developmental acclimation at 15 □°C or 25°C. Sample size varied according to the availability of flies and can be assessed by inspection of the supplementary methods. A single observed scored all flies for all manual runs.

### Time to knock-down

Time to knock-down (TKD) was measured at three noxious constant temperatures. Further, for one of these KD temperatures, we assessed the effect of prior heat hardening. Because *D. melanogaster* and *D. subobscura* have different thermal tolerance we used species specific KD, developmental acclimation, and hardening temperatures. *D. melanogaster* were reared at 25°C and then hardened for 1 hour at 25, 33 or 35°C, whereas *D. subobscura* were reared at 19°C and hardened at 19, 31 or 33°C for 1 hour (followed by recovery at the developmental temperature for 1h), with hardening temperatures corresponding to mild or severe heat hardening, respectively. Hardening was performed in a circulating water bath followed by For *D. melanogaster*, KD temperatures were set to either 35, 37, or 39□°C and for *D. subobscura*, the temperature was set to either 33, 35, or 37□°C corresponding to ‘low’, ‘medium’, or ‘high’ levels of stress for these species. Sample size varied according to the availability of flies and can be assessed by inspection of the supplementary methods.

### Analysis of video recordings

Rather than manually score video recordings, we used Ethovision XT (v10.x) tracking software (Noldus, Wageningen, The Netherlands) to generate activity data over time for each individual (see supplementary methods, Figures S1-4). We note that our method is applicable to any tracking software provided it generates a continuous metric such as distance moved or pixel change per time unit. The raw tracking data (as distance moved, recorded 30 times per second) was exported so that it could be analyzed in R. First, we aggregated the data over time to be the average movement every 15 seconds. Next, we quality filtered the data by looking at the activity profiles generated for each individual. In order to be included in the analysis, flies had to exhibit some decrease in activity over time (indicating a cessation of movement). Visual inspection of the videos showed that excluded individuals (that did not exhibit normal behavior) were either missing or damaged/dead during the set-up (never moved during the video). Finally, in a few cases flies that were not detected by the tracking software were also excluded (see supplementary methods for individual data and account of the removed individuals, Figures S1-4).

Behavioral CTmax or TKD was then determined as the time taken until the activity of flies decreased below a threshold (as values for technical reasons did never reach zero, see supplementary methods, Figure S5). This movement threshold was determined by an algorithm, which identified the maximum activity level of the noisy data after activity progressively declined during assays and was further validated by investigating the effect on tolerance estimates of selecting different thresholds (see supplementary methods, Figures S6-9). The procedure and quality control is presented in Figure 1, and an example of the R-script used to estimate CTmax and TKD is available in the supplementary methods. For CTmax the time taken until cessation of movement was then converted to a temperature based on the measured (i.e. realized) ramping rate (see below). The average treatment specific activity profiles for both CTmax and TKD of both species are available in the supplementary methods (Figures S10-13).

**Figure 1.**
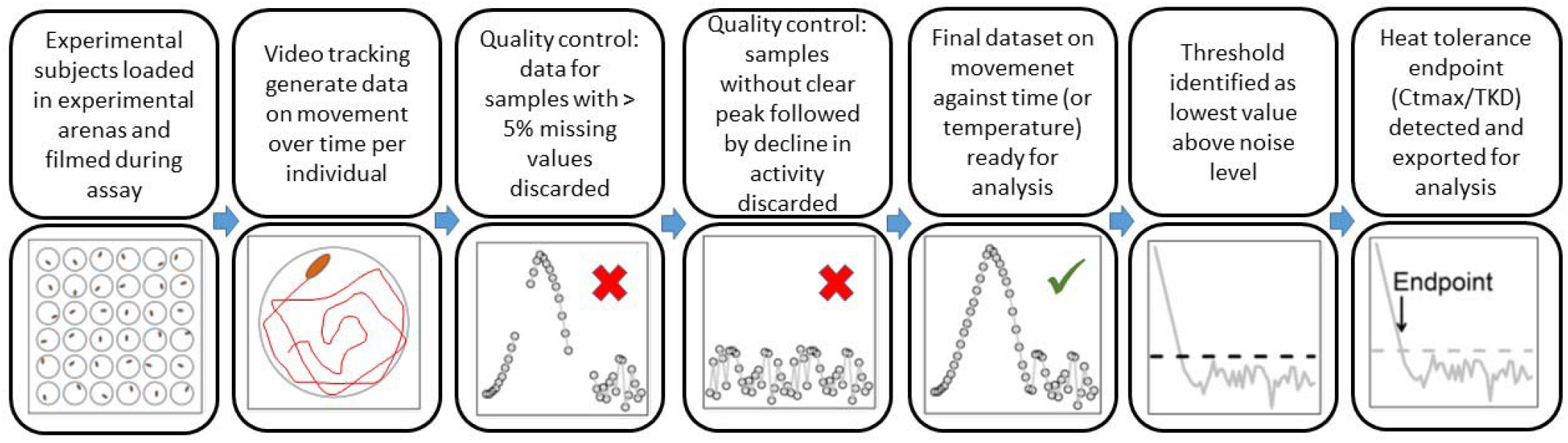
Overview of workflow for automation of thermal tolerance using a combination of motion-tracking software and R.

### Temperature dynamics in water and air

Our manual estimate of CTmax relies on ramping temperature up in water. A previous study showed that realized ramping rates are very close to our programmed rate. The mean realized ramping rate for a programmed rate of 0.1 °C/min across 61 experiments was 0.0984 ± 0.001 (SD) °C/min (Schou et al. 2016). In this study the realized rate of temperature change was calculated from a linear correlation of the measured vial temperatures to 0.49 °C/min (set rate: 0.50 °C/min) and to 0.098 °C/min (set rate 0.10 °C/min). Our automated estimate of CTmax relies on ramping temperature up in air. As the heat transfer is less efficient in air as compared to water, we carefully measured the rate of temperature change inside the cabinet and inside experimental arenas, as these might deviate from each other (see supplementary methods, Figure S14). Temperatures were logged for each experiment and the realized rates (as determined from linear correlations between time and temperature) were reported in all cases and used to convert time to CTmax in temperature. We also determined the time lag in heating the experimental arenas (i.e. the difference in temperature measured in the programmed cabinet and experienced by the experimental subject in the arena at a given time). This time lag corresponded to 0.99 °C for a ramping rate of 0.12 °C/min to 2.54 °C for a ramping rate of 0.47 °C/min (see supplementary methods, Table S1). Thus, CTmax values for both ramping rates used here were calculated from time using the observed (measured ramping rates and starting temperatures (intercepts) of linear correlations) temperatures during the assay (see supplementary methods, Figure S15).

For TKD we also measured the dynamics of temperature change in the experimental arenas of the air cabinet during assays. We performed replicate measurements of set temperatures of 30, 34, 38, and 42 °C. Experimental arena temperature asymptotically approached the set temperature, with the difference decreasing log-linearly. No differences were found between the log-linear decreases in neither slope nor intercept among KD temperatures, suggesting that the temperature dynamics did not differ between low (30 °C) and high (42 °C) KD temperatures (see supplementary methods, Figure S16).

### Statistical analysis

All statistical analysis was carried out using R version 3.6.1 (R Core Team 2019). For dynamic assay estimates, we used linear models for each species and ramping rate to estimate acclimation response between the two treatments. Both were entered as categorical factors in the analyses. For the static assays we also used linear models to estimate the effect of knock-down temperature (continuous factor), and within knock-down temperatures to estimate hardening effect between control and two different hardening temperatures (treated as a categorical factor).

## Results

### CTmax

For *D. melanogaster*, in both the manually scored (physiological) and the automated (behavioural) heat tolerance assays, there was a significant effect of ramping rate and acclimation temperature, with no significant interaction between the two. Effect sizes for developmental acclimation temperature were comparable among the two assay types (model estimate: manual assay 0.13 ± 0.03 °C/°C, automated assay 0.15 ± 0.05 °C/°C). Similar results were found for *D. subobscura*, with the exception that no significant effect was found for developmental acclimation temperature in the manually scored assay. Still, effect sizes for developmental acclimation temperature were similar among the two assay types (model estimate: manual assay −0.03 ± 0.07°C/°C, automated assay −0.04 ± 0.05°C/°C) (Fig. 2, Table 1). As expected CTmax estimates were higher for *D. melanogaster* than for *D. subobscura* in all treatment combinations (mean 3.4°C difference, paired t-test, t_(7)_ = 7.5, p = 0.0001).

**Figure 2.**
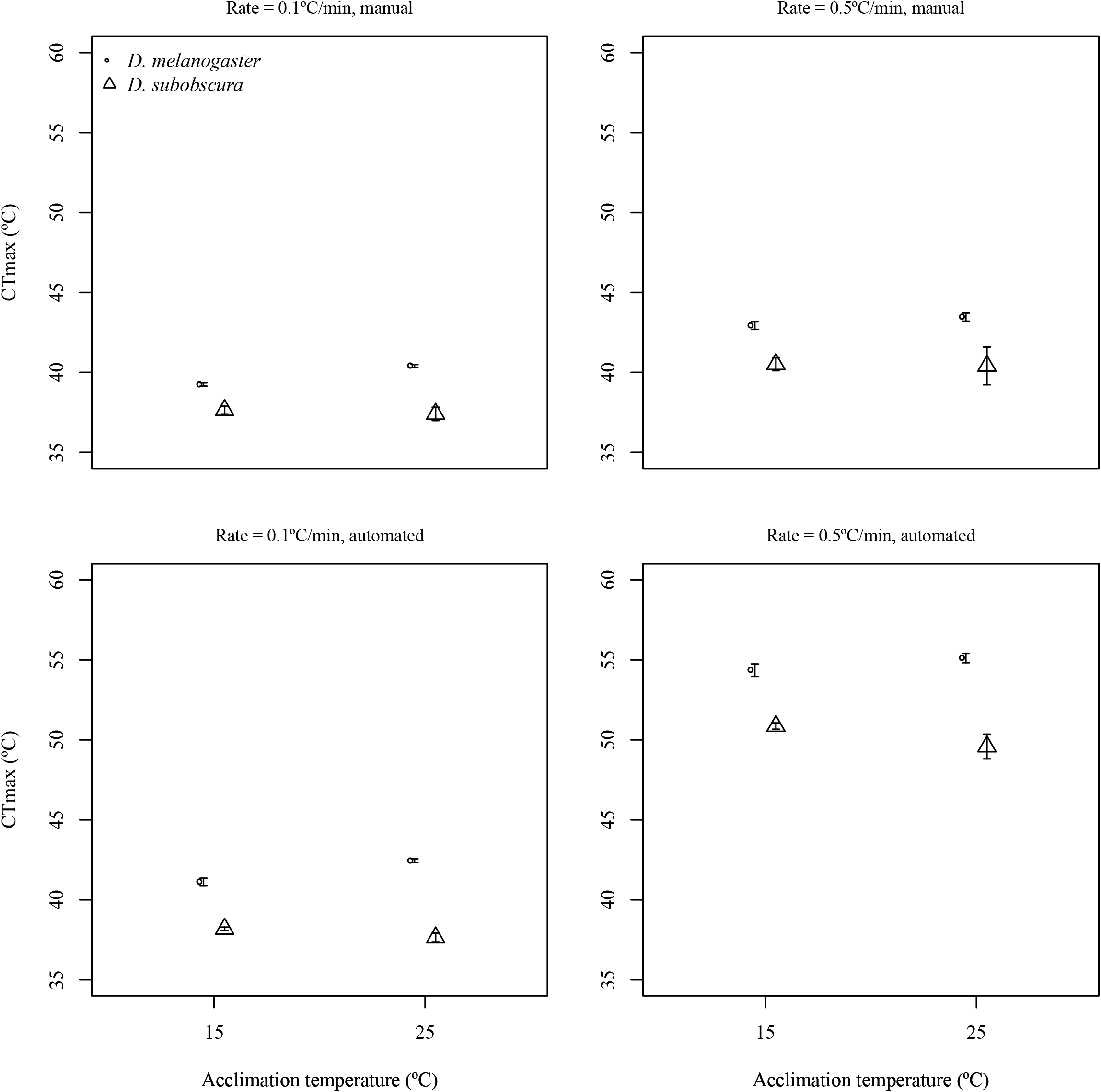
CTmax (mean ± sem) of *D. melanogaster* and *D. subobscura* determined at two ramping rates and after two developmental acclimation temperatures. Top panels show the results from a standard manually scored assay. Bottom panels show the results of the tracking of videos and application of our automated algorithm.

**Table 1.** Effect of ramping rate (0.1 or 0.5 °C/min), developmental acclimation temperature (15 or 25 °C) and their interaction on heat tolerance (CTmax) for two species of *Drosophila* (*D. melanogaster* and *D. subobscura*) as assayed by a manually and an automatically scored assay. Table gives F-values for the ANOVA with degrees of freedom in brackets. *** signifies P < 0.001.

|  | *D. melanogaster* |  | *D. subobscura* |  |
| --- | --- | --- | --- | --- |
|  | Manual | Automated | Manual | Automated |
| Rate | 329.6<sub>(1,80)</sub>*** | 2482<sub>(1,80)</sub>*** | 44.0<sub>(1,73)</sub>*** | 2199<sub>(1,71)</sub>*** |
| Accl | 21.9<sub>(1,80)</sub>*** | 16.4<sub>(1,80)</sub>*** | 0.2<sub>(1,73)</sub> | 8.7<sub>(1,71)</sub>** |
| Rate*Accl | 2.9<sub>(1,80)</sub> | 1.2<sub>(1,80)</sub> | 0.02<sub>(1,73)</sub> | 1.6<sub>(1,71)</sub> |

### Time to Knock-down (TKD)

For both manual (physiological) and automated (behavioural) assays in both species, a non-significant positive effect of the moderate hardening treatment and a significant negative effect of the severe hardening treatment with comparable effect sizes was found (Table 2). As expected, for both species we found a significantly negative relationship between knock-down temperature and knock-down time, in both manual and automated approaches. However, the slopes for the manually scored assays far exceeded the slopes for the automated assay and thus qualitatively differed among assay type (Fig. 2, Table 2). A formal species comparison was not performed for TKD, as developmental acclimation, hardening and KD temperatures differed between the two species. However, both species were tested at a KD temperature of 37°C (corresponding to ‘mild’ for *D. melanogaster* and ‘high’ for *D. subobscura* on Fig. 3). A qualitative comparison clearly shown that TKD estimates were higher for *D. melanogaster* according to our expectation, although with different effect size for the two types of assay (around 60 min for the manual and around 30 min for the automated assay).

**Figure 3.**
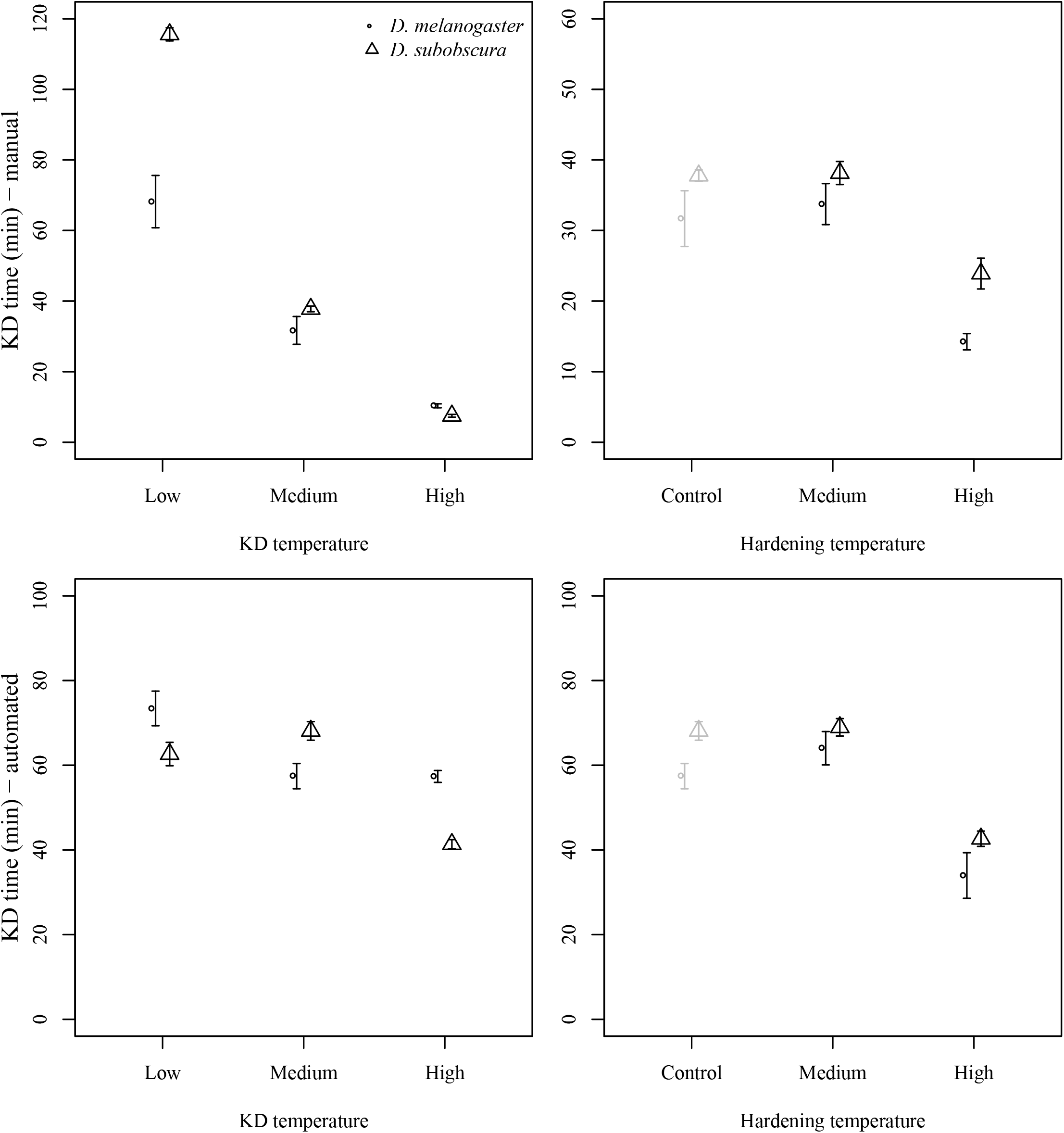
TKD (mean ± sem) of *D. melanogaster* and *D. subobscura* determined at three knock-down temperatures and three hardening temperatures. Top panels show the results from a standard manually scored assay. Bottom panels show the results of the tracking of videos and application of our automated algorithm. KD temperatures were 37, 39 & 41 °C for *D. melanogaster* and 33, 35 and 37 °C for *D. subobscura*, respectively. Pre-treatment (hardening) temperatures were 19 °C as a control, and 37 and 39 °C (for *D. melanogaster*), and 33 and 35 °C (for *D. subobscura*). Note that controls in the right hand side panels (plotted in grey) are the same data as those plotted for the medium KD temperature in the left side panels.

**Table 2.** Effect of KD temperature, and for medium and high temperature hardening for on heat tolerance (time to knock-down) for two species of *Drosophila* (*D. melanogaster* and *D. subobscura*) as assayed by a manually and an automatically scored assay. Table gives F-values for the ANOVA with degrees of freedom in brackets. *** signifies P < 0.001. Model estimates for the effect of KD temperature and for the effect of hardening are in minutes for degree (negative values indicating that KD time shortens as KD temperature is increased).

|  | *D. melanogaster* |  | *D. subobscura* |  |
| --- | --- | --- | --- | --- |
|  | Manual | Automated | Manual | Automated |
| KD temp | 70.5<sub>(1,57)</sub>*** | 13.8<sub>(1,55)</sub>*** | 741.5<sub>(1,58)</sub>*** | 59.8<sub>(1,94)</sub>*** |
| Estimate | -14.5±1.7 | -4.0±1.1 | -27.0±1.0 | -5.4±0.7 |
| Medium hard | 0.2<sub>(1,36)</sub> | 1.7<sub>(1,33)</sub> | 0.04<sub>(1,38)</sub> | 0.1<sub>(1,49)</sub> |
| Estimate | 0.1±0.3 | 0.4±0.3 | 0.03±0.15 | 0.1±0.2 |
| High hard | 18.8<sub>(1,37)</sub>*** | 16.7<sub>(1,27)</sub>*** | 35.7<sub>(1,38)</sub>*** | 86.9<sub>(1,37)</sub>*** |
| Estimate | -1.0±0.2 | -1.3±0.3 | -1.0±0.2 | -1.8±0.2 |

## Discussion

Methodological approaches to generate phenotypic data have not “caught-up” to those of high-throughput molecular data, and thus are a limiting step for many experimental studies (Kong, Axford et al. 2016). The generation of phenotypic data typically rely on manually scoring individuals and exercising observer judgement which can be, at best difficult to compare and at worst, fraught with observer biases (Holman, Head et al. 2015). Here we present a case-study of automation using two *Drosophila* species. We took a two-pronged approach to validate our technique of measuring critical thermal limits: 1) we used species with well-characterized thermal biology, and 2) we directly compare to the manually scored methods across both static and dynamic thermal assays. While, the idea is similar to previous methods suggested for aquatic or terrestrial ectotherms (Burton, Zeis et al. 2018, Awde, Fowler et al. 2020), we are the first to validate the automated method by comparing UTLs across several treatments using manually scored assays. Furthermore, we verified the generation of activity profiles during heat assays needed for automated analysis on a number of different taxa including rather docile organisms such as woodlice and millipedes (see Fig. S17 and S18). Thus, we show that the method described is applicable to arthropods generally. We considered automation successful if the method captures CTmax and TKD estimates similar to those of traditional assays, known species differences and effect sizes of pre-treatments and treatment, is high-throughput, and/or reduces variance attributable to observer imprecision.

We find that automation generally produces qualitatively comparable species differences, and comparable effect sizes for acclimation and hardening responses, while not necessarily producing the same numeric values as our manually scored assay. While the absolute CTmax estimates are not directly comparable, the species and treatment differences (effect sizes) were very similar to the manual assay. This demonstrates that the automation is reliably capturing behavioral upper thermal limits. We note here that we are the first to test automation of these data against the traditional assays. The discrepancy between assay types was most pronounced for the higher CTmax estimates, e.g. at the high ramping rate. It was expected that higher ramping rates leads to higher CTmax, likely due to the inflated accumulation of heat damage at longer assays (low rates) (Sørensen, Loeschcke et al. 2013, Salachan, Burgaud et al. 2019). However, while the automated assay at the low ramping rate exceeded that of the manual assay by only 2-3°C, the automated CTmax estimate of the fast rate exceeded that of the manual by at least 10°C. It is not clear why the difference between the assay types were so strongly affected by ramping rate, however, we propose that it might be a combined contribution from both technical and biological effects. These results highlight the potential bias of interpretation and comparison absolute CTL values generated in different ways without validation.

The manual assay produced a sharper linear decline in TKD estimates with increasing temperatures, corroborating studies across several species of *Drosophila* (Castaneda, Rezende et al. 2015, Jørgensen, Malte et al. 2019, Salachan, Burgaud et al. 2019), compared to automated assay. In the manual assays, for both CTmax and TKD, the endpoint is scored as the cessation of all movement following stimulation (physical tapping the vials and shining a bright light at the flies) which likely corresponds to death or near death of the fly (Lutterschmidt and Hutchison 1997). In contrast, the automated assays necessarily score the endpoint as the cessation of voluntary movement, which might be different from stimulated movement ability (MacLean, Kristensen et al. 2017). Notably, flies in experimental arenas quite quickly regained movement after removal from the thermal cabinets in the TKD experiments. It seems that flies in the automated assay (when exposed to constant high temperatures) stop voluntary movement after a time period not very different among the different KD temperatures used here, suggesting the behavior is an integral of the trait scored from the videos. While clearly different from the physiological capacity as measured by the manual assay, it could be equally important for ecologically relevant heat tolerance estimation as behavior is assumed to be important response to adverse environmental conditions (Sunday, Bates et al. 2014, Logan, van Berkel et al. 2019). In the CTmax experiments, we did not observe a similar difference, and expect that the difference between voluntary and stimulated movement was diminished as temperature continue to increase toward physiological collapse. Thus, while absolute values should not be compared to the values of traditional assays, we propose that the automated CTmax assay does capture a large proportion of the physiological component of heat tolerance, similarly to the manually scored assay. This is supported by very similar effect size estimates of hardening responses across the two assay types.

We find that our method provides several additional benefits and only a single drawback with respect to absolute values. First, it has been suggested that variation among individuals within treatment groups in different assays can be inflated by the inability of the observer to accurately monitor all individuals in real time (Castaneda, Calabria et al. 2012). We investigated this potential intra-observer measurement error by comparing automated to manual scores across treatments. We found that the automation resulted in only marginally decreased standard deviations for CTmax (F = 4.04_(1,14)_, P = 0.064), and no effect on TKD estimates (F= 0.002_(1,18)_, P = 0.97). In all but one case, the effect of acclimation for *D. subobscura* in CTmax (Table 1), these minor differences did not lead to qualitatively different interpretation of the results. Thus, there are no clear indications that the manual assays suffer from inflated observer induced variance (Castaneda, Calabria et al. 2012). Second, the number of subjects that can be handled by a minimal set-up of our method is 72 (two arenas filmed simultaneously), requiring one light-board, one camera, and one thermal cabinet). With additional arenas prepared during assays (as they do not require attention), a single researcher can process hundreds of subjects per day. This can easily be up-scaled, depending on access to equipment leading to much larger throughput. While we used a commercial tracking software (Ethovision XT, Noldus, Wageningen, The Netherlands), our method does not rely on this particular software (see e.g. Awde, Fowler et al. 2020) and can be applied to any software that can generate data on distance per unit time. Third, in addition to the endpoint used (cessation of movement), the collected data contain other potentially interesting information. The plots of average activity profiles (Figures S10-13) showed treatment specific patterns, indicating that the maximal movement or temperature of maximal movement could reveal biological relevant information about the effect of acclimation, ramping rate or species differences. This needs to be explored further in dedicated studies, but might mean that individual thermal tolerance can be estimated before physiological collapse is reached. This might have important implications for studies applying selection or estimating heritability (e.g. Hangartner and Hoffmann 2016).

Several earlier studies have used manual scoring of video recordings (Woods and Bonnecaze 2006, Hazell, Pedersen et al. 2008, MacLean, Higgins et al. 2016, Awde, Fowler et al. 2020). However, none of the previous studies validated their assays against manually scored assays. Further, while manual scoring of video recordings allow the observers to repeat scoring for decreased measurement error, it does not automate the process. The automated approach described here proved to be robust to experimental conditions, treatments, and selection of threshold (see Fig. 1). Individual arenas exclude the potential effect of density and interaction among individuals (Hazell, Pedersen et al. 2008). We conclude that the automated method generates unbiased and reproducible results and can be used in experiments with large number of samples. We find that effect sizes of acclimation and hardening responses are determined as accurately as for the traditional methods, but that the endpoint measured in the automated method might be different and incorporate a potentially overlooked and ecologically relevant behavioral aspect of thermal tolerance (Sunday, Bates et al. 2014). Finally, we find no evidence of the assays scored in real time suffer from inflated variance and thus are not flawed by intra-observer inaccuracy or imprecision as suggested elsewhere (Rezende, Tejedo et al. 2011, Castaneda, Calabria et al. 2012). Thus, we suggest that our automated method is beneficial in terms of through-put, but that both methods are generating biological meaningful results. However, further investigations should elucidate the discrepancy of absolute values among assays. Furthermore, automated scoring might be applied to scoring of cold tolerance, entry and recovery from chill coma (Hazell, Pedersen et al. 2008) and to other tedious laboratory traits (in many arthropod species), such as desiccation or starvation tolerance, with similar benefits to through-put. As such trait are usually scored at predefined intervals, rather than continuously, automated assays can potentially improve resolution markedly.

## Acknowledgements

We thank Trine Bech Søgaard and Annemarie Højmark for excellent help in the fly lab and Paul Vinu Salachan for helpful discussions and comments to an earlier draft of the manuscript. This work was supported by a grant from Aarhus University Research Foundation to JGS.

